# *Rare but not absent*: the Inverted Mitogenomes of Deep-Sea Hatchetfish

**DOI:** 10.1101/2023.06.12.544378

**Authors:** André Gomes-dos-Santos, Nair Vilas-Arrondo, André M. Machado, Esther Román-Marcote, Jose Luís Del Río Iglesias, Francisco Baldó, Montse Pérez, Miguel M. Fonseca, L. Filipe C. Castro, Elsa Froufe

**Affiliations:** CIIMAR/CIMAR - Interdisciplinary Centre of Marine and Environmental Research, University of Porto, Matosinhos, Portugal; Programa de Doctorado “Ciencias marinas, Tecnología y Gestión” (Do*MAR), Montse Pérez Facultad de biología, Universidad de Vigo, Vigo, Spain; Centro Oceanográfico de Vigo Instituto Español de Oceanografía (IEO, CSIC). Subida a Radio Faro, 50. 36390 Vigo (Pontevedra), Spain; Centro Oceanográfico de Cádiz, Instituto Español de Oceanografía (IEO, CSIC), Puerto pesquero, dársena de Levante s/n, 11006 Cádiz, Spain; Department of Biology, Faculty of Sciences, University of Porto, Porto, Portugal

**Author notes:** Corresponding authors: Elsa Froufe and L. Filipe C. Castro.

**Keywords:** mitogenome, strand asymmetry, deep-sea fish, vertebrate, mitochondrial gene order, rearrangements

## Abstract

Mitochondrial genomes are by definition compact and structurally stable over aeons. This generalized perception results from a vertebrate-centric vision, as very few types of mtDNA rearrangements have been described in vertebrates. By combining a panel of sequencing approaches, including short- and long-reads, we show that species from a group of illusive marine teleosts, the deep-sea hatchetfish (Stomiiforms: Sternoptychidae), display a myriad of new mtDNA structural arrangements. We show a never reported inversion of the coding direction of protein-coding genes (PGG) coupled with a strand asymmetry nucleotide composition reversal directly related to the strand location of the Control Region (which includes the heavy strand replication origin). An analysis of the 4-fold redundant sites of the PCGs, in thousands of vertebrate mtDNAs, revealed the rarity of this phenomenon, only found in 9 fish species, five of which are deep-sea hatchetfish. Curiously, in Antarctic notothenioid fishes (Trematominae), where a single PCG inversion (the only other record in fish) is coupled with the inversion of the Control Region, the standard asymmetry is disrupted for the remaining PCG but not yet reversed, suggesting a transitory state in this species mtDNA. Together, our findings hint that a relaxation of the classic vertebrate mitochondrial structural *stasis*, observed in Sternoptychidae and Trematominae, promotes disruption of the natural balance of asymmetry of the mtDNA. Our findings support the long-lasting hypothesis that replication is the main molecular mechanism promoting the strand-specific compositional bias of this unique and indispensable molecule.

## Introduction

Counterintuitively, the mitochondrial genome (mtDNA) is far more variable than normally recognized, including structure, gene content, order and orientation, organization and mode of expression (Shtolz and Mishmar 2023). In Metazoa, several types of mtDNA are described, including linear telomeric molecules (e.g., Cnidaria), “giant” circular molecules (e.g. some Bivalvia and Placozoa) and several mini-circular molecules (e.g. Insecta) (Kolesnikov and Gerasimov 2012; Shtolz and Mishmar 2023). In vertebrates, the classic mtDNA is described as a compact circular molecule, maternally inherited, between 16–19 kbp and two compositionally distinct strands, conventionally referred to as heavy (H-strand with high G composition) and light (L-strand with low G composition) (Clayton 1982; Boore 1999; Kolesnikov and Gerasimov 2012). The standard vertebrate mtDNA encodes 37 genes, 13 protein-coding genes (PCG); two ribosomal RNAs (rRNAs); 22 transfer RNAs (tRNAs); and two differently located and strand-specific replication origins (heavy strand replication origin [OH] and light strand replication origin [OL]) (Clayton 1982; Lee et al. 1995; Boore 1999; Kolesnikov and Gerasimov 2012). The OH is placed within a larger non-coding region, the Control Region (CR), which includes several regulators of the mtDNA replication and transcription and where four evolutionary conserved sequence blocks are commonly found (i.e, CSB-I, CSB-II, CSB-III, and CSB-D) (Clayton 1982; Lee et al. 1995; Boore 1999; Kolesnikov and Gerasimov 2012). The 13 mtDNA PCG, which follows the standard architecture (i.e., number, relative positioning and coding strand), play key functional roles in the oxidative phosphorylation (OXPHOS) cascade (Clayton 1982; Boore 1999; Kolesnikov and Gerasimov 2012).

Exceptions to this generalized picture, in vertebrate classes, are extremely rare, with very few and small-scale deviations (Gissi et al. 2010; Zhang et al. 2020; Formenti et al. 2021; Montaña-Lozano et al. 2022; Sharbrough et al. 2023; Shtolz and Mishmar 2023). Among the different groups, birds and reptiles show the highest distribution of rearranged genes, while mammals and fish show residual examples of rearrangements (Zhang et al. 2020; Montaña-Lozano et al. 2022; Shtolz and Mishmar 2023). In fish, three distinct types of mtDNA gene rearrangements have been described recently by Satoh et al. (2016): “shuffling”, i.e., local position change maintaining coding polarity (i.e., genes coding strand); “translocation”, i.e., movement to a distinct location maintaining the genes encoded strand; and the rarest event “inversions”, i.e., genes (or non-coding unities) switch to their complementary strand (Fonseca et al. 2008; Kong et al. 2009; Gong et al. 2013; Fonseca et al. 2014; Arrondo et al. 2020; Papetti et al. 2021; Shtolz and Mishmar 2023). Inversions, when acting upon the replication controlling unities (i.e., the CR or OH), have been shown to promote a switch of the strand asymmetry nucleotide composition. Consequently, The mtDNA replication is suggested to be involved in the differently accumulated strand mutation patterns of the mtDNA (assuming an asymmetric model of mtDNA replication) (Fonseca et al. 2008; Fonseca et al. 2014; Shtolz and Mishmar 2023). Marine deep-sea hatchetfish from the family Sternoptychidae (Order: Stomiiformes) are a group of small (less than 100 mm) and peculiar mesopelagic fishes (Nelson 2016). The family consists of two subfamilies, Maurolicinae and Sternoptychinae, which include around 70 species distributed through 10 genera (Howell and Krueger 1987; Nelson 2016; Coad 2019). These species are generally found in high abundance and biomass within the mesopelagic ichthyofauna and with important ecological functions and a key trophic position (Gjoesaeter and Kawaguchi 1980; Eduardo et al. 2020). Species from the family Sternoptychidae have been described as “*some of the most bizarre stomiiforms*” and are characterized by having a condensed body with a reflecting fattened silver side that allows camouflage (Carnevale 2008). As in all other Stomiiformes, deep-sea hatchetfish possess specialized bioluminescent organs, i.e., photophores that allow them to produce light (Krönström et al. 2005; Carnevale 2008; Haddock et al. 2009).

Here we describe four new Sternoptychidae whole mtDNA, using Illumina PE short reads and Oxford Nanopore long reads, showing that the mtDNA of Sternoptychidae have a diverse and unusual gene architecture. Our findings include mtDNA gene shuffling, translocation and inversions, acting on PCG, tRNA rRNA and/or CR. In particular, we demonstrate that strand asymmetry nucleotide composition reversal occurs when genes change their coding polarity relative to the Control Region (i.e. OH, initiation of replication). Conversely, inversions of CR were shown to promote the complete nucleotide strand asymmetry reversal in two deep-sea hatchetfish species. Moreover, by investigating over 6000 species we determine that strand asymmetry is a rare event in vertebrate mitochondrial genomes. In Antarctic notothenioid fishes (Trematominae), a PCG inversion (the only case previously reported in fish) coupled with the CR inversion is shown to disrupt the standard asymmetry for the remaining PCG. Together, our findings provide strong evidence supporting the long-lasting theory that replication is the main molecular mechanism promoting the strand-specific compositional bias of the mtDNA.

## Results and Discussion

### Deep-sea hatchetfish mtDNA show a widespread complex repetitive region hampering full sequence circularization

We generated four novel whole mitochondrial genomes from the marine hatchetfish, two of them resulting from new nuclear genome assemblies: *Argyropelecus aculeatus* Valenciennes, 1850 (lovely hatchetfish), *Argyropelecus hemigymnus* Cocco, 1829 (half-naked hatchetfish), *Argyropelecus olfersii* (Cuvier, 1829) and *Maurolicus muelleri* (Gmelin, 1789) (Mueller’s pearlside). The raw sequencing reads and mtDNA assemblies were deposited at NCBI, and respective SRA and assembly accessions are depicted in Table 1 and linked to BioProject PRJNA977192 (Provisory Reviewer Link https://dataview.ncbi.nlm.nih.gov/object/PRJNA977192?reviewer=qp1erbs33ac430jhnvu9demc1m).

**Table 1.** List of whole mitogenomes used in the whole mitogenome-based phylogeny. * represent mitogenomes sequenced in this study.

| Family | Species | GenBank Accession | BioSample Accession | SRA Accession | mtDNA length (bp) |
| --- | --- | --- | --- | --- | --- |
| Sternoptychidae | *Argyropelecus aculeatus* * | OR062951 | SAMN35453525 | SRR24764805 and SRR24764804 | 23,291 |
| Sternoptychidae | *Argyropelecus affinis* | AP012964.1 | - | - | 15,489 |
| Sternoptychidae | *Argyropelecus hemigymnus* * | OR062952 | SAMN35453526 | SRR24764802 | 20,747 |
| Sternoptychidae | *Argyropelecus olfersii* * | OR062953 and OR062954 | SAMN35453527 | SRR24764803 | 16,414 |
| Sternoptychidae | *Maurolicus muelleri* | AP012963.1 | - | - | 15,230 |
| Sternoptychidae | *Maurolicus muelleri* * | OR062955 | SAMN35453528 | SAMN35453528 | 16,604 |
| Sternoptychidae | *Polyipnus polli* | AP012962.1 | - | SRR24764801 | 16,773 |
| Sternoptychidae | *Sternoptyx diaphana* | MT588184.1 | SAMN35453678 | SRR24764797-SRR24764800 | 17,224 |
| Sternoptychidae | *Sternoptyx obscura* | OP057081.1 | - | - | 18,293 |
| Gonostomatidae | *Sigmops gracilis* | AB016274.1 | - | - | 16,436 |
| Stomiidae | *Tactostoma macropus* | LC377784.2 | - | - | 17,563 |
| Stomiidae | *Photonectes margarita* | AP018417.1 | - | - | 18,592 |
| Stomiidae | *Trigonolampa miriceps* | AP012961.1 | - | - | 15,709 |
| Stomiidae | *Astronesthes lucifer* | AP012959.1 | - | - | 15,491 |
| Stomiidae | *Stomias atriventer* | MG321595.1 | - | - | 17,596 |
| Stomiidae | *Chauliodus sloani* | AP002915.1 | - | - | 17,814 |
| Diplophidae | *Diplophos taenia* | AP012960.1 | - | - | 16,427 |
| Diplophidae | *Diplophos sp.* | AB034825.1 | - | - | 16,418 |
| Phosichthyidae | *Vinciguerria nimbaria* | AP006769.1 | - | - | 16,741 |
| Phosichthyidae | *Vinciguerria nimbaria* | AP012958.1 | - | - | 16,741 |
| Myctophum affine | *Myctophum affine* | AP002922.1 | - | - | 16,239 |
| Trachipteridae | *Trachipterus trachipterus* | AP002925.1 | - | - | 16,162 |
| Ateleopodidae | *Ateleopus japonicus* | AP002916.1 | - | - | 16,650 |
| Synodontidae | *Saurida undosquamis* | AP002920.1 | - | - | 15,737 |
| Salmonidae | *Coregonus lavaretus* | AB034824.1 | - | - | 16,737 |

The mtDNA lengths varied from 15,230 bp in *M. muelleri* to 23,291 bp in *A. aculeatus*, largely influenced by poorly resolved repetitive regions, which includes the putative CR and long species-specific intergenic regions (figs 1-3). The only previously assembled hatchet fish mtDNA with a CR annotation is *Polyipnus polli* Schultz, 1961 (AP012962.1) (fig. 1). Conversely, in other publicly available mtDNAs, the CR was either not annotated, i.e., in *Sternoptyx obscura* Garman, 1899 (OP057081.1) and in *Sternoptyx diaphana* Hermann, 1781 (MT588184.1) (fig. 2), or not sequenced, i.e., in *M. muelleri* (AP012963.1) and *Argyropelecus affinis* Garman, 1899 (AP012964.1) (fig 1). The difficulty in obtaining this region has already been observed in many other animal groups (e.g., (Ghiselli et al. 2021)). Conversely, in the three novel *Argyropelecus sp.* sequenced mtDNAs, i.e., *A. aculeatus*, *A. hemigymnus*, and *A. olfersii*, long intergenic repeats were identified (fig. 3). The difficulty in resolving these repetitive regions using short reads prevented the assembly of a single contig in *A. olfersii* (composed of two contigs), as well as the circularization of the *A. hemigymnus* assembly (fig. 3). This factor also hampered the assembly of the recently published *S. diaphana* mtDNA, which required scaffolding with Sanger sequencing (Arrondo et al. 2020). Nevertheless, we efficiently identified the CSB-II (thus the CR) in all but one (i.e., *A. olfersii*) (Supplementary File 1 and fig. S1).

**FIG.1.**
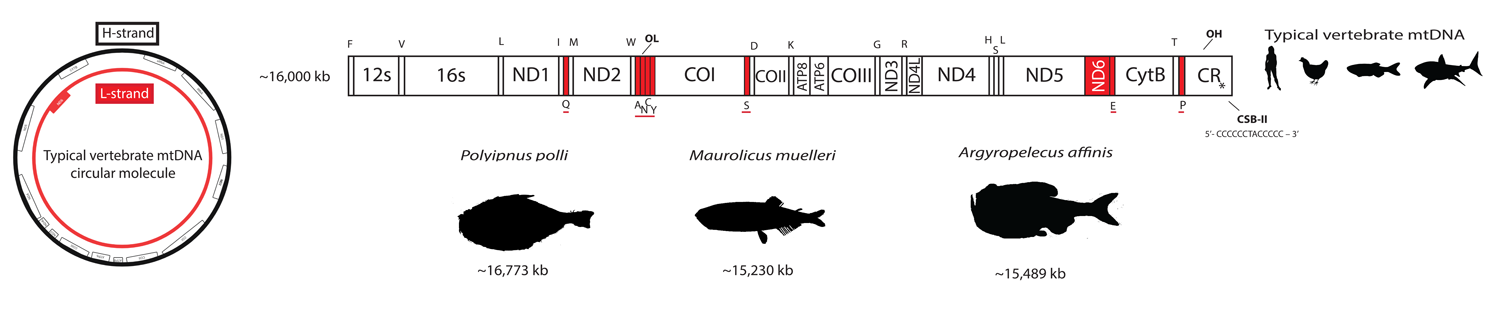
Left: Schematic representation of circular mitochondrial molecule highlighting the protein-coding genes encoded in the different strands. Right: Representation of linearized mitochondrial molecule, depicting the standard vertebrate gene order shared by the deep-sea hatchetfish species, *Polyipnus polli*, *Maurolicus muelleri* and *Argyropelecus affinis*. Genes encoded in the L-strand are depicted in red; Genes encoded in the H-strand are depicted in white. * Control Region has not been sequenced in either *Maurolicus muelleri* or *Argyropelecus affinis*. The CSB-II sequence is a representation of the generally expected composition of the motif.

Sequencing approaches are key for resolving complex mtDNA assemblies (e.g. (Calcino et al. 2020; Formenti et al. 2021)). Consequently, in *A. aculeatus* we used Oxford Nanopore long-reads to determine the composition of the complete mtDNA molecule. The produced reads were highly efficient in resolving the full mtDNA sequence, with some reads even spawning the entire molecule, supporting the inferred architecture. The final assembly was circularized, revealing two long repetitive regions, one spawning ∼4800 bp, which included the CSB-II (between tRNA-Asp and tRNA-Pro) and the other ∼1750 bp (between tRNA-Phe and a COI) (figs 3 and S2). As for the other two *Argyropelecus sp.*, although the short-read assemblies retrieved all the mtDNA genes, the aforementioned repetitive regions were fragmented (fig. 3).

### Gene shuffling, translocations, duplications and inversions define the structure of mtDNA structure in deep-sea hatchetfish

We next investigated the gene content and overall structure of the mtDNA in deep-sea hatchetfish. The newly sequenced genomes as well as those deposited in GenBank show striking deviations from the standard vertebrate gene arrangement (figs 1-3). While the mtDNA from three species, i.e., *M. muelleri, A. affinis* and *P. polli*, maintain the standard vertebrate mtDNA architecture (fig. 2), most of the newly sequenced mtDNA revealed considerably modified architectures (figs 2-3) (Saccone et al. 2002; Satoh et al. 2016). The results show that shuffling, translocation and inversion of PCGs, rRNAs and more abundantly tRNAs, are widespread in the mtDNA deep-sea hatchetfish (figs 2-3). Even within the same genus, high structural differences can be detected (figs 2-3). Species from the *Argyropelecus* genus show four distinct mtDNA architectures: one maintains the standard vertebrate structure; while the other three share a radical inversion of a large fragment composed of several genes, with reciprocal additional unique features (fig. 3). This fragment includes the inversion of the PCGs ND2 and ND1, the rRNA 16S and 12S and several tRNAs (M, I, L1, V and F), as well as the shuffling of two tRNAs (A and C) (fig. 3). The inversion of two PCGs has never been reported in fish mtDNA before. To date, the only reported inversion of a PCG is of ND1 in a single clade of Antarctic notothenioid fishes (Nototheniidae: Trematominae) (Papetti et al. 2021; Minhas et al. 2023). Interestingly, this also resulted from the inversion of a large fragment, which included the rRNA genes, several tRNAs (E, I, L2, V and F) as well as the CR (Papetti et al. 2021; Minhas et al. 2023).

**FIG.2.**
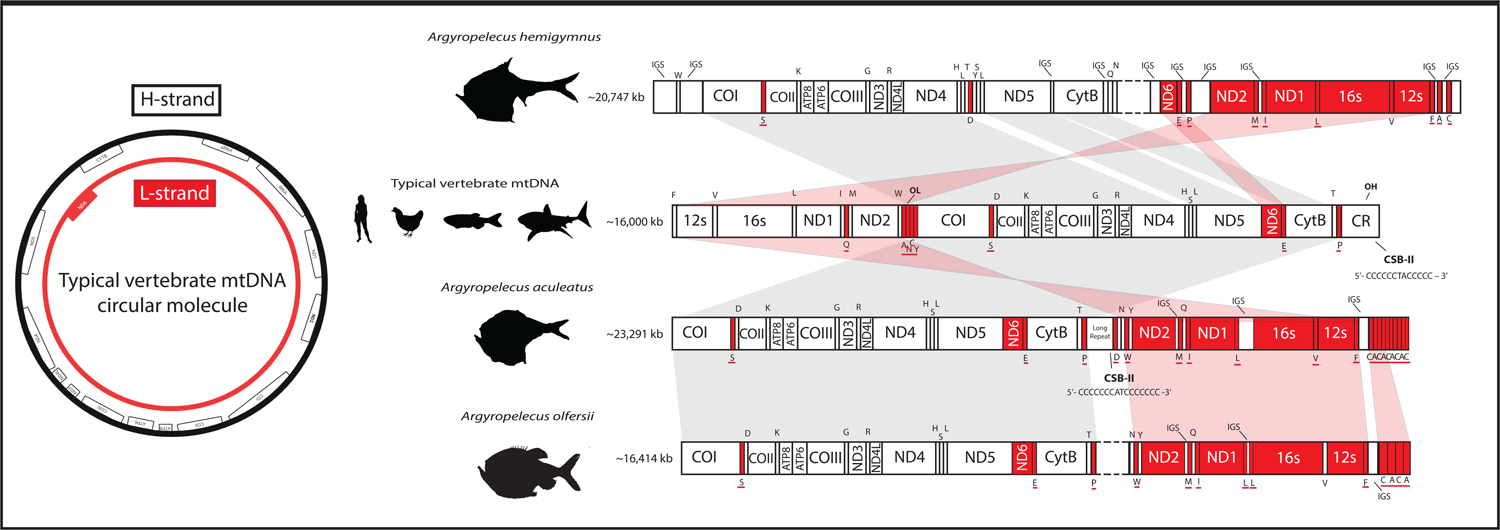
Left: Schematic representation of circular mitochondrial molecule highlighting the protein-coding genes encoded in the different strands. Right: Representation of linearized mitochondrial molecule, depicting the standard vertebrate gene order (middle) comparatively to deep-sea hatchetfish species, *Sternoptyx obscura*, *Sternoptyx diaphana*. Genes encoded in the L-strand are depicted in red; Genes encoded in the H-strand are depicted in white; Red shadow represents macrosyntenic patterns of gene inversion. The CSB-II sequence for the typical vertebrate mtDNA is a representation of the generally expected composition of the motif. The CSB-II sequences for the marine hatchet fish are the ones here identified.

**FIG.3.**
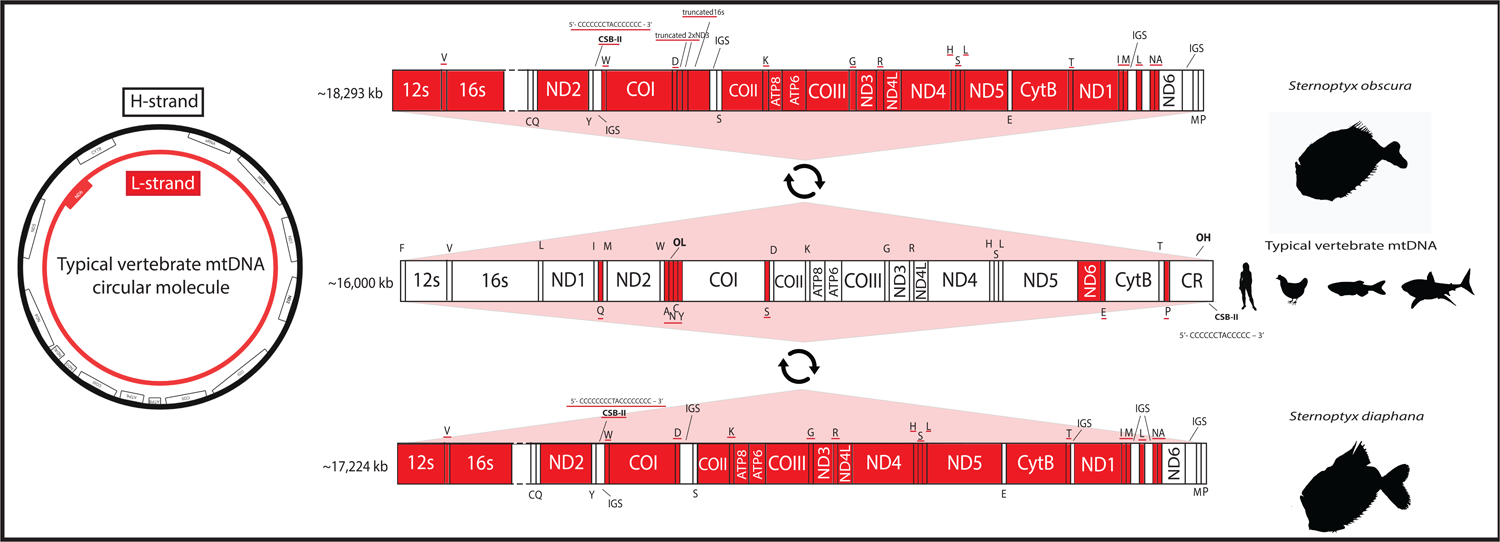
Left: Schematic representation of circular mitochondrial molecule highlighting the protein-coding genes encoded in the different strands. Right: Representation of linearized mitochondrial molecule, depicting the standard vertebrate gene order (second from the top) comparatively to deep-sea hatchetfish species, *Argyropelecus hemingymnus*, *Argyropelecus aculeatus* and *Argyropelecus olfersii*. Genes encoded in the L-strand are depicted in red; Genes encoded in the H-strand are depicted in white; Red shadow represents macrosyntenic patterns of gene inversion; Grey shadow represents macrosyntenic patterns of genes in the same strand. The CSB-II sequence for the typical vertebrate mtDNA is a representation of the generally expected composition of the motif. The CSB-II sequences for the marine hatchet fish are the ones here identified.

Some species-specific tRNA duplications were observed (figs. 2-3). The most striking of these duplications was captured by the Nanopore-based assembly in *A. aculeatus*, corresponding to a tandem duplication of two tRNAs C-A, followed by one tRNA C. This pattern seems to also occur in *A. olfersii*, in which a shorter tandem duplication of two tRNAs C-A was present, while in *A. hemigymnus* the two tRNAs are present in the same location but in a single copy (fig. 3). Furthermore, *A. hemigymnus* shows a unique shuffling of the ND6 and the tRNA E (fig. 2). The two *Sternoptyx sp.* Revealed the overall same mtDNA architecture, with *S. obscura* having duplications and “*pseudogenization*” of ND3 and 16S (fig. 2). Gene duplication in mtDNA is often followed by the loss of one the copies, frequently retaining fragments of the original duplicated gene (Satoh et al. 2016; Papetti et al. 2021).

### Phylogenetic Analysis shows poorly resolved deep-sea hatchetfish evolutionary relationships

We next analysed the phylogenetic relationships between the various deep-sea hatchetfish species. The resulting phylogenetic inference, rooted with *Coregonus lavaretus* (Linnaeus, 1758) (following (Ijichi et al. 2018; Arrondo et al. 2020)), is split into two major groups, one composed of order Stomiiformes and the other composed of families Synodontidae, Ateleopodidae, Myctophidae and Trachipteridae (fig. 4). Stomiiformes’ phylogenetic relationships are poorly revolved, with very low support in most of the nodes. Within deep-sea hatchet fish, i.e., family Sternoptychidae, the relationships among the genus are also poorly resolved and an unexpected long branch is seen for the two S*ternoptyx* species (fig. 4). The low support persisted in the amino acid-based phylogeny (fig. S3). The mtDNA phylogeny here presented is the first to include more than one species of the deep-sea hatchet fish (see (Arrondo et al. 2020)). However, the taxon representation is still incomplete, which likely explains the low support observed. The relationships of the four *Argyropelecus sp.* are well resolved and reflect the structural mtDNA variation here described (figs. 1-3). *Argyropelecus affinis*, the only species to preserve the standard vertebrate structure is the first split within the genus, followed by *A. hemigymnus*, sister to *A. olfersii* + *A. aculeatus*, the two most structurally similar (fig. 4).

**FIG.4.**
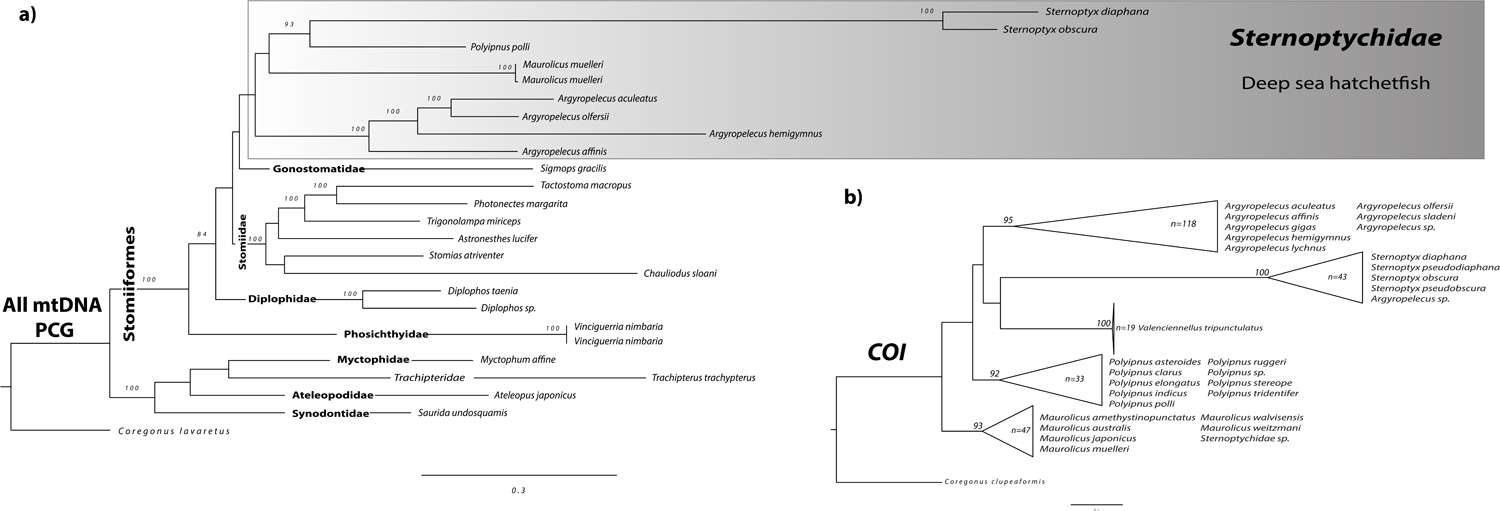
a) IQ-tree Maximum Likelihood (ML) phylogenetic inference retrieved from the nucleotide alignment of the 13 mitogenomes concatenated protein-coding genes. b) IQ-tree Maximum Likelihood (ML) phylogenetic inference retrieved from the nucleotide alignment of all available COI sequences of deep-sea hatchetfish in NCBI. Above the nodes are represented ultrafast bootstrap values above 80%.

To obtain a more taxa-representative phylogeny, all the cytochrome oxidase subunit I (COI) sequences available on GenBank were used (fig. 3). The COI alignment included a total of 261 sequences (total length of 652 bp) (Supplementary File 2), from 26 distinct deep-sea hatchetfish species and the outgroup species, i.e., *Coregonus clupeaformis* (Mitchill, 1818). The resulting nucleotide phylogeny shows once again low support among inter-genus relationships (fig. 4). Even though this phylogeny includes 260 sequences from marine hatchetfish, it highlights the need to increase the molecular data for the whole group.

### GC/AT skew of strand-specific 4-fold redundant sites show a strikingly frequent strand asymmetry reversal in deep-sea hatchetfish always coupled with CR strand relative position

The mtDNA asymmetric strand nucleotide composition of the H and L strands is generally highly demarked in vertebrates, with a strong signature in the A-T and C-G composition of each stand (Saccone et al. 2002). This pronounced signature is the distinguishable factor between the two strands, i.e., the H(heavy)-strand, which is guanine rich, and L(light)-strand, which is guanine poor (Francino and Ochman 1997). Conversely, inverted genes, i.e., genes that change to the complementary strand, will be exposed to the mutational bias specific of the new strand, and thus are expected to change their mutational signature accordingly, as proposed by the asymmetric model of mtDNA replication (Fonseca et al. 2008; Kennedy et al. 2013; Fonseca et al. 2014).

Given the newly detected inversions of two PCGs in *A. aculeatus*, *A. hemigymnus*, and *A. olfersii*, as well as the oddly long branch lengths observed in *Sternoptyx sp.*, we next estimated the GC/AT skew of the PCGs at the 4-fold redundant sites (the most likely to accumulate strand-specific mutations and to reflect the underlying mutational processes given that selection acting on these sites is weaker) for all the marine hatchetfish under study.

As expected, in the mtDNA that maintains the standard vertebrate architecture with no detectable structural changes, i.e., *M. muelleri*, *A. affinis* and *P. polli*, shows the AT/GC skew pattern of a gene encoded in the L-strand (fig. 5 and S4, i.e., AT skew < 0 and GC skew > 0). Interestingly, in the two *Argyropelecus sp.* with inverted PCG polarity, this AT/GC skew pattern was observed in ND6, but also in ND1 and ND2, the two PCGs included in the inverted fragment (fig. 5 and S4). Most strikingly, for the two *Sternoptyx sp.*, the entire AT/GC skew pattern is reversed in all PCGs (fig. 5 and S4) but with no rearrangements or inversions detected in those genes. What could be causing the AT/GC skew reversion in these genomes? To date, such phenomenon, in vertebrates, has been suggested to result from CR inversion, and from the replication mechanism itself with the leading strand becoming the lagging strand and vice-versa during an asymmetric mode of replication (Fonseca et al. 2008; Fonseca et al. 2014). We tested this hypothesis by characterising the relative positioning of the initiation of replication and by identifying elements that have been described to be key in the replication mechanism of vertebrate mtDNA: the conserved sequence blocks CSB-II and III. Typically, CSB-II consists of six or more cytosines, one thymine and one adenine ending with five or more cytosines: 5’-CCCCCCTACCCCC – 3’. We used this sequence as a reference, allowing for some variation in the length of the poly-C flanks. CSB-III is less conserved, generally identified by its positioning regarding CSB-II, and may be absent in fish CR (Satoh et al. 2016). The Motif Discovery analysis identified CSB-II in all the analysed sequences, except in *P. polli* while CSB-III was less prevalent, as previously observed by Satoh et al., (2016) (Supplementary File 1 and fig. S1). The absence of both CSBs in *P. polli* needs further evaluation, as this mitogenome is a direct submission to NCBI without detailed information on how it was generated.

**FIG.5.**
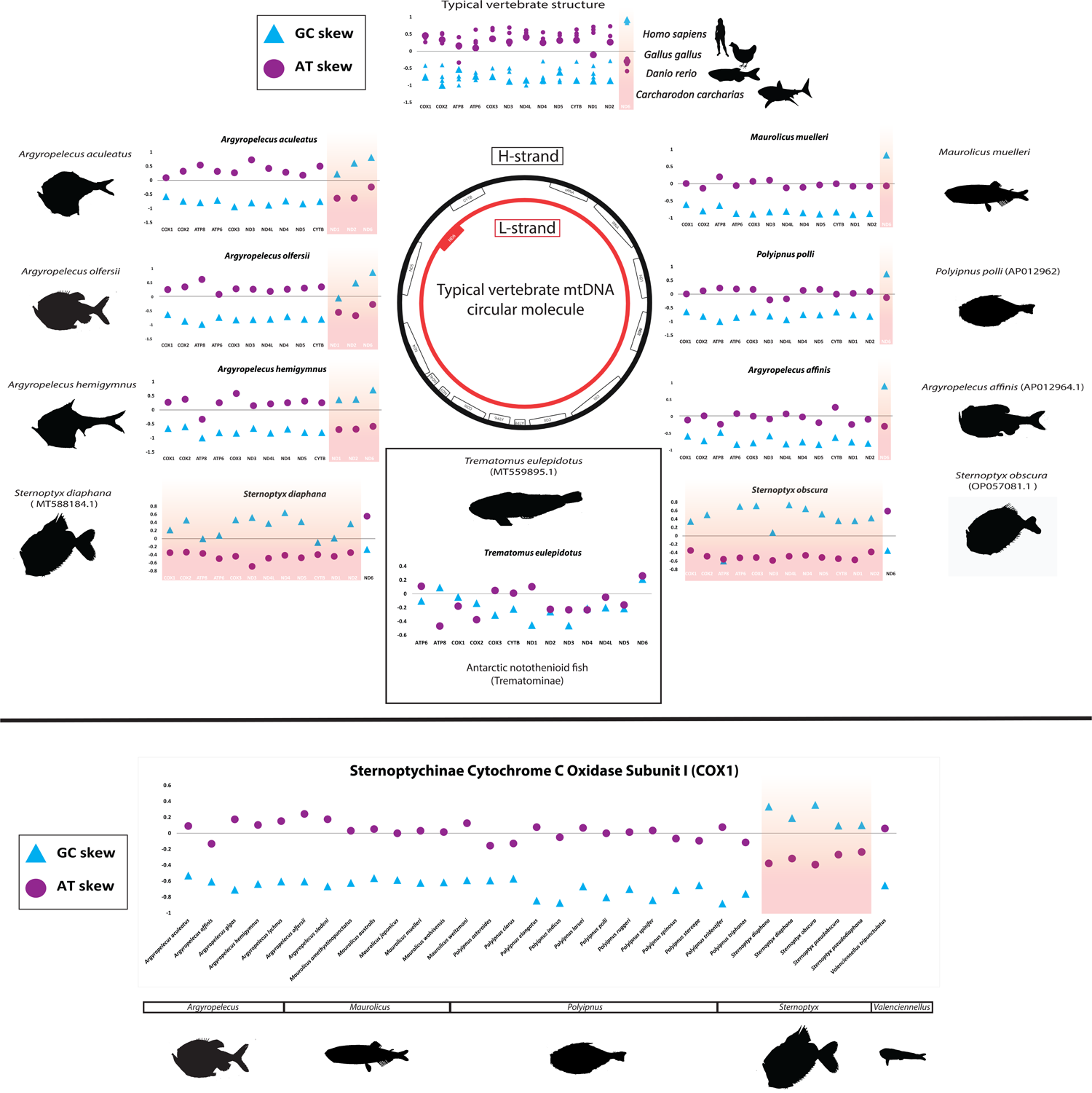
Top: Strand-specific 4-fold redundant sites GC/AT skew estimations for all protein-coding genes, for four model vertebrate species with standard vertebrate mitogenome gene order, each species of deep-sea hatchetfish analysed and one representative species from the Antarctic fish subfamily Trematominae that have an inversion of the ND1 and CR (inside the box). Bottom: Strand-specific 4-fold redundant sites GC/AT skew estimations for COI mitochondrial gene of all deep-sea hatchetfish, available in NCBI. Red shadows highlight the inversion of the strand-specific nucleotide GC/AT skew pattern. Middle: Schematic representation of circular mitochondrial molecule highlighting the protein-coding genes encoded in the different strands.

Astonishingly, as previously predicted, the orientation of the CSB-II and therefore of the CR relative to the PCG was the common denominator determining the AT/GC skew nucleotide asymmetry pattern (Supplementary File 1 and figs 1-3 and S1). That is, in the two *Sternoptyx sp.* the CSB-II is the only element changing its coding polarity (inversion), having the same polarity as ND6 and contrary to all other PCGs, while in the remaining species, the relative position follows the standard vertebrate architecture (Supplementary File 1 and figs 1-3 and S1).

To date, the only reported PCG inversion in a vertebrate is in one clade of Antarctic notothenioid fishes (Nototheniidae: Trematominae), consisting of the inversion of a large genomic segment (∼5kbp), which includes four tRNAs, two rRNAs, the PCG inversion (ND1) and the CR (Kennedy et al. 2013; Papetti et al. 2021). Once again, the relative position of the CR, here identified through CSB-II (Supplementary File 1 and fig. S1), is the determining factor shaping the nucleotide strand asymmetry, i.e., ND1 is the only PCG with the same polarity as the CR and thus it is also the only PCG maintaining its AT/GC skew pattern, while the remaining genes show a completely disrupted pattern. Interestingly, the fact that the latter-mentioned PCG shows a disruption but not a reversal AT/GC skew pattern, suggests that, in this group of organisms, the process of the mtDNA nucleotide composition strand asymmetry reversal is still in a transitory state (fig. S5). A complete disruption of the typical strand asymmetry signature was detected in all 15 Trematominae mtDNA available, while the remaining notothenioid fishes maintained the standard pattern (fig. S5).

To access similar putative hidden patterns in other vertebrates, we next performed a vertebrate-wide assessment of the AT/GC skew at 4-fold redundant sites (Fonseca et al. 2008; Fonseca et al. 2014) in a total of 6,297 mtDNA (Retrieved from RefSeq in May 2022). The results demonstrate that the complete reversal of the nucleotide strand asymmetry composition is observed solely in the *Sternoptyx sp.* and those previously reported by Fonseca et al., (2008, 2014), totalizing seven species (figs. 6 and S4). Moreover, single gene asymmetry reversal is detectable for the ND1 and ND2 genes of the three *Argyropelecus* species here described (figs. 6 and S4). Given that there are still many un-sequenced marine hatchetfish whole mtDNA, we hypothesised that calculating the AT/GC skew pattern at 4-fold redundant sites for the most sequenced mitochondrial gene of the group, i.e., COI, would be an informative strategy regarding the whole family. The results also show the reversed pattern in all analysed *Sternoptyx* species (n=4), thus suggesting the reversal of the nucleotide strand asymmetry is shared by all analysed *Sternoptyx* (figs. 6 and S4). The remaining hatchetfish species maintain the standard vertebrate AT/GC skew pattern, which is also in accordance with the results deduced from whole mtDNA assembly and annotation.

### Genomic *Oddities*: inverted mtDNA and large nuclear genomes

The combination of sequencing methodologies allowed the determination of *de novo* four mtDNA from deep-sea hatchetfish species. Strikingly, the sequential analysis of gene annotation, phylogenetics, and AT/GC skew sequence investigations support a unique case of mtDNA structural plasticity (Shtolz and Mishmar 2023). We expanded the analysis to include thousands of available mtDNA. Our findings show that partial or full mtDNA inversions are evolutionary restricted in a minute of unrelated fish clades: *Argyropelecus sp*. *Sternoptyx sp*. The immediate causes of this phylogenetically restricted alteration of mtDNA structure are unknown and deserve further exploration. In fact, within Metazoa, vertebrates show the strongest signs of purifying selection for gene rearrangement (Shtolz and Mishmar 2023), with recent studies suggesting that gene blocks inversions (only recorded in invertebrates) promote sticking changes in the transcriptional pattern, thus requiring new regulatory elements (Blumberg et al. 2014; Blumberg et al. 2017; Barshad et al. 2018; Shtolz and Mishmar 2023). Consequently, our highly unexpected findings raise the interesting possibility of an adaptive scenario of the reported mtDNA oddities. Interestingly, using the high-coverage whole genome sequencing for *A. aculeatus* and *S. diaphana* we were able to appraise the genome size for both species (fig. S6). The results show an unexpectedly large genome size estimation, with both species being approximately 2.70 Gbp long (fig. S6). Most teleost species have an average of 1 Gbp long genome size, with the exceptions being usually attributed to whole genome duplication events (Parey et al. 2022). Together, our results suggest that like the mitogenomes, the whole genome of deep-sea hatchetfish is also unusual and deserves future explorations.

## Methods

### Samples collection and DNA extraction

In total, four specimens were captured for this study, three from the genus *Argyropelecus*, i.e., *Argyropelecus aculeatus* Valenciennes, 1850, *Argyropelecus hemigymnus* Cocco, 1829, and *Argyropelecus olfersii* (Cuvier, 1829), and one from genus *Maurolicus*, i.e., *Maurolicus muelleri* (Gmelin, 1789). The specimens were obtained during the scientific surveys: EU Groundfish Survey (Platuxa-2019) in North Atlantic (43,1544 N-51,4429 W, 2019); from the EU Groundfish Survey (Platuxa-2019) in the Northwest Atlantic Ocean (43,3838 N-49,0036 W, 2019); from the Survey PORCUPINE20 In Porcupine Bank (51.0677 N; −14.2862 W, 2020) and from EU Groundfish Survey FN3L19 in North Atlantic (−47,668 N; 47,497833 W), respectively. Morphological identification was performed onboard and whole specimens are stored in absolute ethanol and are stored at DNA and Tissue bank at CIIMAR – Interdisciplinary Center of Marine and Environmental Research. The specimen treatment has been approved by the CIIMAR ethical committee and by CIIMAR Managing Animal Welfare Body (ORBEA) according to the European Union Directive 2010/63/EU. Whole genomic DNA for each specimen was obtained from a small portion of the muscle tissue using the Qiagen MagAttract HMW DNA extraction kit, following the manufacturer’s instructions. For all samples, total DNA was used for Illumina paired-end (PE) library preparation and sequencing at the Macrogen, Inc. (Seoul, Korea), using Illumina HiSeq X Ten platform, with 150 bp PE configuration. For *A. aculeatus* and *S. diaphana* high coverage whole genome (WGS) PE sequencing was performed, while the remaining samples were only sequenced at low coverage. Furthermore, low-coverage Nanopore (Oxford Nanopore) genome skimming was performed for *A. aculeatus*. Briefly, ∼1 µg of genomic DNA was used for library preparation using the LSK109 kit and after sequenced on an FLO-MIN106 revD SpotON R9.4 Flow Cell for 48 h.

### Whole mitogenome assemblies and annotation

Raw Illumina PE reads were quality-filter and adaptors were removed using Trimmomatic (version 0.38) (Bolger et al. 2014), using the parameters LEADING:5 TRAILING:5 SLIDINGWINDOW:5:20 MINLEN:36. Read quality was inspected before and after trimming using FastQC (version 0.11.8) (http://www.bioinformatics.babraham.ac.uk/projects/fastqc/). Whole mitogenome reconstruction for each species was obtained using several distinct approaches. For each species, assemblies were performed with GetOrganelle v1.7.1 (Jin et al. 2020), specifying a multi k-mer approach (i.e., from 20-125 with a 5-mer increment). GetOrganelle is an interactive baiting pipeline that filters mtDNA reads and uses the SPAdes assembler (Bankevich et al. 2012) to reconstruct the mitogenome with the filtered reads. The results of the assemblies were individually validated using multiple approaches. For cases where multiple assemblies were generated, i.e., *A. hemigymnus*, and *A. olfersii*, the gfa files were inspected using Bandage v0.8.1 (Wick et al. 2015) which revealed several ambiguous disjoints. Furthermore, annotation was generated for some of the putative assemblies using the module “annotate” from MitoZ v.3.4 (Meng et al. 2019), which showed that the ambiguous disjoints were localized within no-coding repetitive regions. Consequently, new assemblies were performed, first using larger k-mer with GetOrganelle v1.7.1 and if the problem persisted, using metaSPAdes v3.12.0 (Nurk et al. 2017) with the maximum K-mer size possible, i.e., 127. In the end, the selected best representative assemblies for each species were as follows, for *M. muelleri* the GetOrganelle assembly with multi k-mer (20-125-mer with a 5-mer increment), for *A. olfersii* the GetOrganelle assembly single k-mer (131-mer) and for *A. hemigymnus* the metaSPAdes assembly with the maximum allowed k-mer size (127-mer). Every generated assembly was annotated with MitoZ (as described above). Read coverage distributions were analysed by aligning PE reads to the final genome assemblies using the Burrows-Wheeler Aligner (BWA) v.0.7.17-r1198 (Li 2013) and visualized in Artemis v17.0.1 (Carver et al. 2012) (fig. S7).

The Nanopore reads of *A. aculeatus* were quality-filtered using Filtlong (https://github.com/rrwick/Filtlong). Given that long repetitive regions seem to cause problems with the PE-based assemblies, the Nanopore reads were filtered by size (i.e., >21,000bp) to include only reads spanning the whole mitogenome. The mitogenome assembly was performed using Unicycler v.0.4.8. (Wick et al. 2017)). The assembly was polished, following the author’s suggestions, with the Nanopore reads using medaka v1.2.2 (https://github.com/nanoporetech/medaka) and with short-reads, first using Polypolish v0.4.3 (Wick and Holt 2022) and after using Polca (Zimin and Salzberg 2020). Genome annotation was performed with MitoZ (as described above). Read coverage distributions were analysed by aligning PE reads (as described above).

Finally, since the *Sternoptyx obscura* mitogenome available on NCBI (OP057081) is marked as “UNVERIFIED” and therefore no annotation is provided, the mitogenome was downloaded and annotated using MitoZ (as described above).

To identify the putative Control Region (CR) of the deep sea hatchet fish mitogenome assemblies, we search for conserved motifs within non-coding regions to identify any of the known Conserved Sequence Blocks (CSB) that are involved in the replication initiation (Satoh et al. 2016). The recently described CSBs of several fish species were here used as a reference to guide the search, including one Stomifformes, i.e. *Diplophos taenia* (Satoh et al. 2016). We focused on the two most conserved of the three CSBs, i.e., CSB-II and CSB-III. Given the lack of significant non-coding regions within the mitogenomes of *M.muelleri*, *A. affinis*, and *A. olfersii* (see Results and Discussion), these mitogenomes were not included in the analysis. Moreover, given the highly disproportionate read coverage distribution in the mitogenome non-coding regions of *A. hemigymnus* (fig. S7), this mitogenome was not considered. The coverage distribution indicates regions likely represented by the collapse of a highly repetitive region, thus not suited for the analysis. The non-coding regions from the remaining deep-sea hatchet fish species, the Antarctic Trematominae species, *Pagothenia borchgrevinkias* (with the only other record CDS mitochondrial inversion), as well as all the fish CSB-II and CSB-III identified by Satoh et al. (2016) were used. The sequences were uploaded to MEME v.5.5.2 (Bailey and Elkan 1994) and the Motif Discovery model was used to identify conserved motifs across all sequences, specifying a maximum motif width of 25.

### Phylogenetic reconstruction

To produce a phylogenetic reconstruction, the whole mitogenomes of all Stomiiformes available on NCBI (n=15), including four Sternoptychidae species (i.e., *S. obscura, M. muelleri*, *P. polli*, *A. affinis*), the mitogenomes here produced and four outgroup taxa were used (Table 1). Alignments of the 13 protein-coding genes (PCGs) were produced using MAFFT v7.453 (Katoh and Standley 2013). Positions with gaps in 50% or more of the individual alignments were removed using trimAL v. 1.2rev59 (Capella-Gutiérrez et al. 2009) and all alignments were concatenated using FASconCAT-G (https://github.com/PatrickKueck/FASconCAT-G). The alignment composed by the concatenated PCGs from 15 species had a total length of 11,436 bp. The partition scheme and the evolutionary models that best fit those schemes, as well as Maximum Likelihood (ML) phylogenetic inference were produced in IQ-TREE v.1.6.12 (Nguyen et al. 2015; Kalyaanamoorthy et al. 2017).

For the amino acid phylogenetic reconstruction, individual alignments were translated to proteins using the EMBOSS seqret V.6.6.0.0, trimmed with trimAL v. 1.2rev59 (as described above) and concatenated using FASconCAT-G. The alignment composed by the concatenated PCGs from 15 species had a total length of 3,801 aa. The partition scheme and the evolutionary models that best fit those schemes, as well as Maximum Likelihood (ML) phylogenetic inference were produced in IQ-TREE v.1.6.12.

### COI Phylogeny

To produce a phylogenetic reconstruction with a wider taxa representation, all COI sequences available on GenBank (n=261) were downloaded (including the outgroup taxa *C. clupeaformis*). Alignment was performed using MAFFT v7.453 (Katoh and Standley 2013) and an ML phylogenetic inference was constructed in IQ-TREE v.1.6.12, also estimating the best evolutionary model for the analysis.

### Strand-specific 4-fold redundant sites GC/AT skew estimations

The protein-coding genes of the complete mitogenomes of all vertebrates available in GenBank (http://www.ncbi.nlm.nih.gov) in May of 2022 were retrieved. The calculation of the GC skew (G-C)/(G+C) and the AT skew (A-T)/(A+T) at 4-fold redundant sites using custom Perl scripts (Fonseca et al. 2014). The redundant codons examined were alanine (GCN), proline (CCN), serine (TCN), threonine (ACN), arginine (CGN), glycine (GGN), leucine (CTN), and valine (GTN). Furthermore, to infer GC/AT skew inversions in other species from family Sternoptychidae, for which no whole mitogenome is available, the COI sequences for all Sternoptychidae species available on GenBank were downloaded and the calculations applied, as described above.

### Whole Genome size estimation

The high coverage WGS PE sequencing reads for *A. aculeatus* and *S. diaphana* were used to estimate the overall characteristics of each species’ genomes using Jellyfish v.2.2.10 and GenomeScope2 (Ranallo-Benavidez et al. 2020) with a k-mers length of 21.

#### Data Availability Statement

The raw sequencing reads and mtDNA assemblies are deposited at NCBI, and respective SRA and assembly accessions are depicted in Table 1, all linked to BioProject PRJNA977192.

FIG.S1. Schematic representation of the MEME - Motif discovery results, including the inferred conserved motif representing CSB-II (in red) and, in some species, CSB-III (in blue). The results include several CR from fish species studied in Satoh et al (2016) (using the same abbreviated nomenclature), as well as the non-coding regions of the deep-sea marine hatchet fish species *Sternoptyx obscura*, *Sternoptyx diaphana*, *Argyropelecus aculeatus*, the Stomiiformes species *Diplophos taenia,* and the Antarctic Trematominae species *Pagothenia borchgrevinki*.

FIG.S2. Complete circular mitochondrial molecule of the deep-sea hatchetfish species *Argyropelecus aculeatus*, produced by CGView/Proksee (https://proksee.ca).

FIG.S3. IQ-tree Maximum Likelihood (ML) phylogenetic inference retrieved from the amino acid alignments of the 13 concatenated mitogenome protein-coding genes.

Above the nodes are represented ultrafast bootstrap values.

FIG.S4. Plots of the GC and AT skews at the 4-fold redundant sites of the mitochondrial protein-coding genes ND3, ND4, ND4L, ND5, COX2, COX3, CYTB, ATP6 ATP8 from 6,297 vertebrate species. Dots represent each of the 6,297 analysed vertebrate species, with red (deep-sea hatchetfish) and green (other fish species) dots highlighting genes with inverted AT and GC skews. The plots for the remaining protein-coding genes are provided in fig.6.

**FIG.6.**
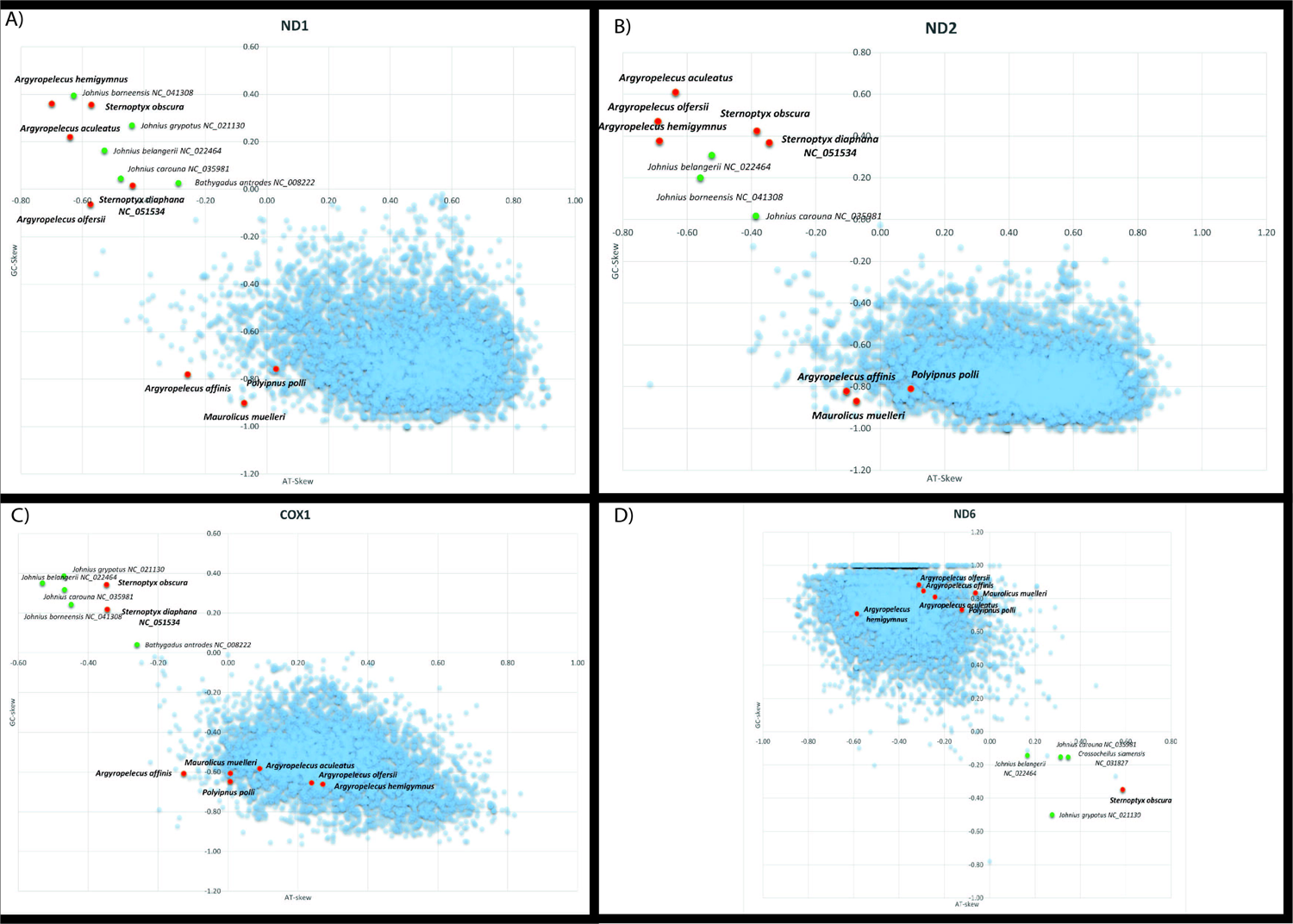
Plots of the GC and AT skews at the 4-fold redundant sites of the mitochondrial protein-coding genes ND1, ND2, COX1 and ND6 from 6,297 vertebrate species. Dots represent each of the 6,297 analysed vertebrate species, with red (deep-sea hatchetfish) and green (other fish species) dots highlighting genes with inverted AT and GC skews. The plots for the remaining protein-coding genes are provided in fig.S4.

FIG.S5. Strand-specific 4-fold redundant sites GC/AT skew estimations for all protein-coding genes, for all mitogenomes of notothenioid fishes available on NCBI. Coloured circles represent the mitochondrial gene orders described by Papetti et al. 2021. Species not studied by Papetti et al. 2021 are in the black box. Mitogenomes with an inversion of the ND1 gene are highlighted in a thick red box.

FIG.S6. GenomeScope2 k-mer (21) distributions displaying estimation of genome size (len), homozygosity (aa), heterozygosity (ab), mean k-mer coverage for heterozygous bases (kcov), read error rate (err), the average rate of read duplications (dup), k-mer size used on the run (k:), and ploidy (p:) for *Argyropelecus aculeatus* (left) and *Sternoptyx diaphana* (right).

FIG.S7. Artemis read coverage plot distribution across the assembled marine hatchet fish mitogenome assemblies.

Supplementary File 1 - MEME - Motif discovery results, including the sequences of the inferred conserved motif representing CSB-II and, in some species, CSB-III. The results include all the CR of the fish species studied in Satoh et al (2016) (using the same abbreviated nomenclature), as well as the non-coding regions deep-sea marine hatchet fish species *Sternoptyx obscura*, *Sternoptyx diaphana*, *Argyropelecus aculeatus*, the Stomiiformes species *Diplophos taenia,* and the Antarctic Trematominae species *Pagothenia borchgrevinki*.

Supplementary File 2 - COI alignment including 261 sequences from 26 distinct deep-sea marine hatchetfish species and the outgroup species, i.e., *Coregonus clupeaformis* (Mitchill, 1818) (total length of 652 bp).

