## Supplementary figures and images for "*Rare but not absent*: the Inverted Mitogenomes of Deep-Sea Hatchetfish"

### Fig.S1.jpg

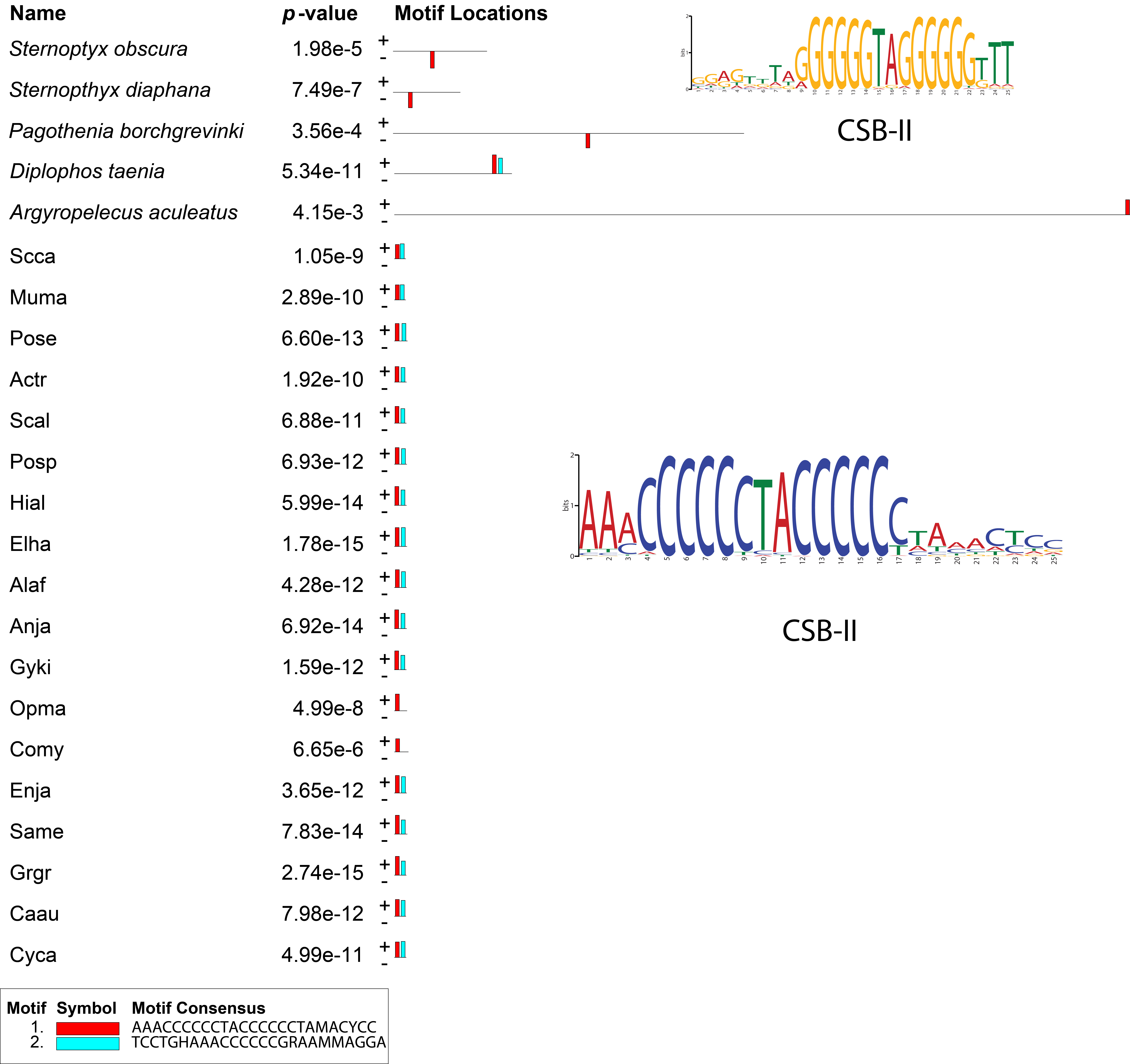

### Fig.S2.jpg

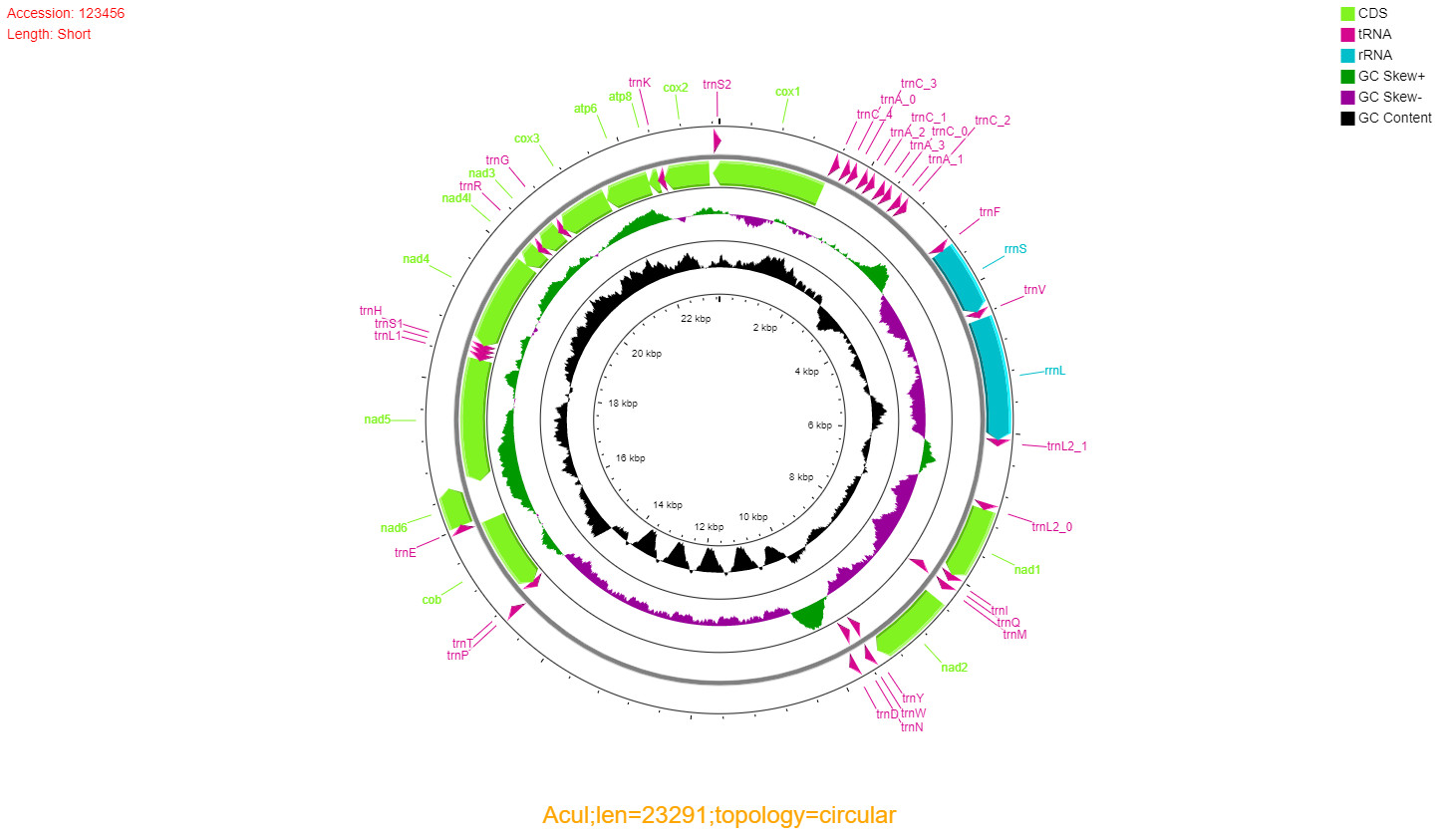

### Fig.S2.png

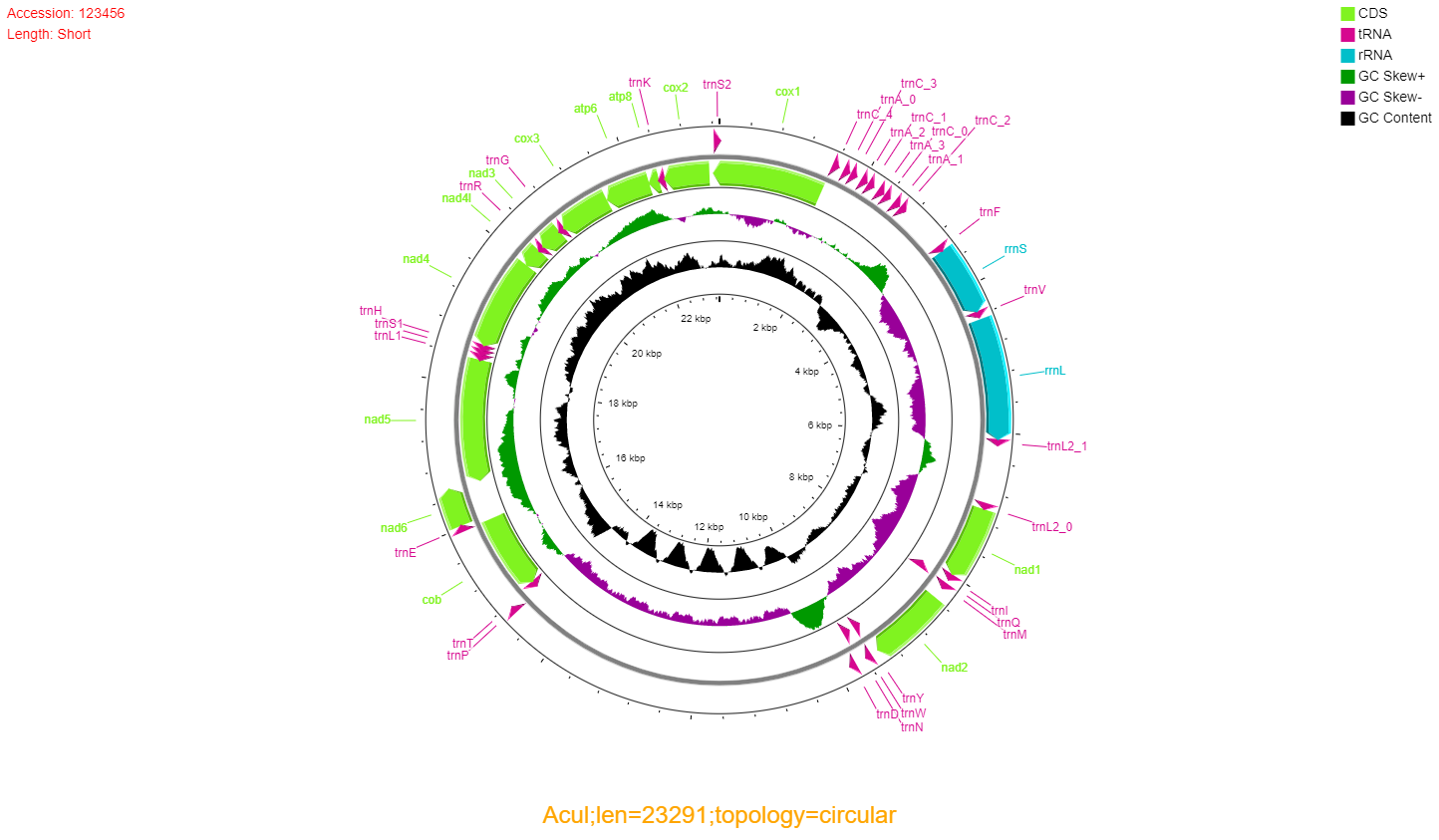

### Fig.S3.pdf

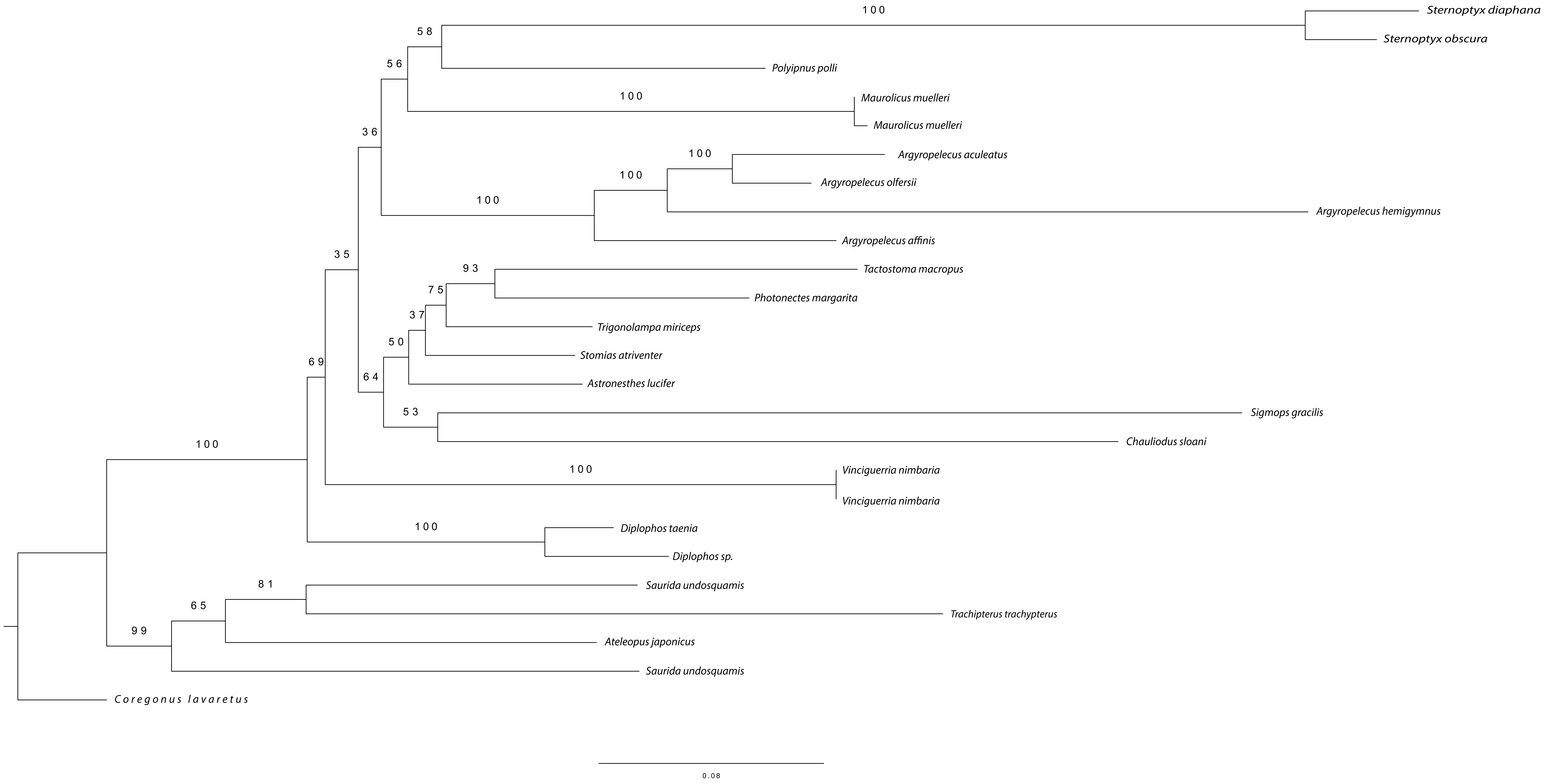

### Fig.S4.jpg

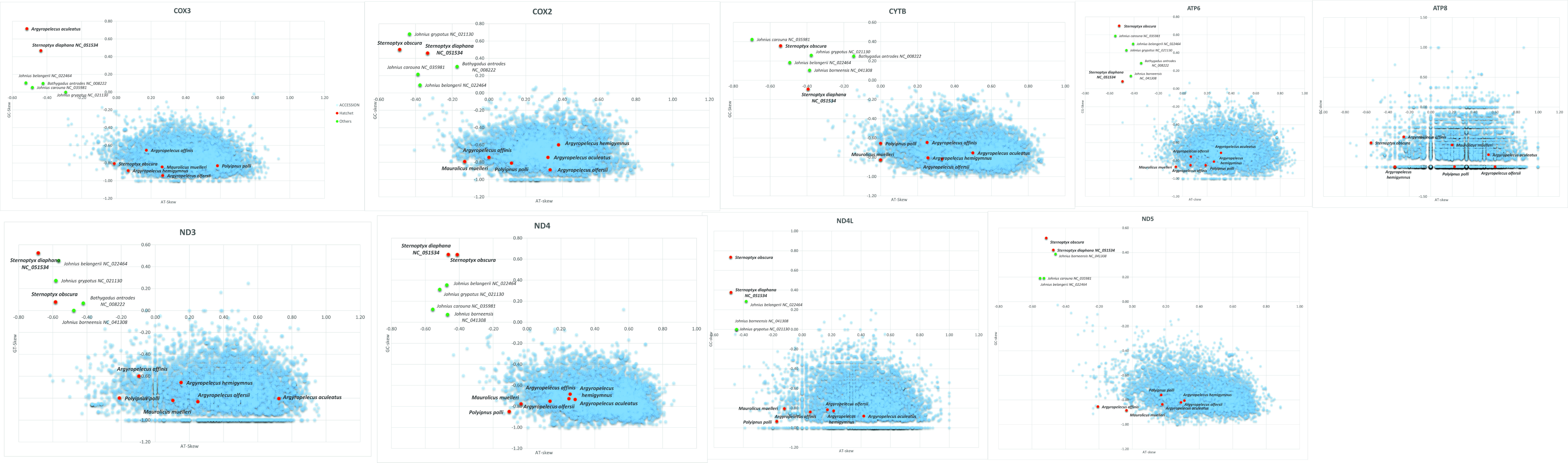

### Fig.S5.pdf

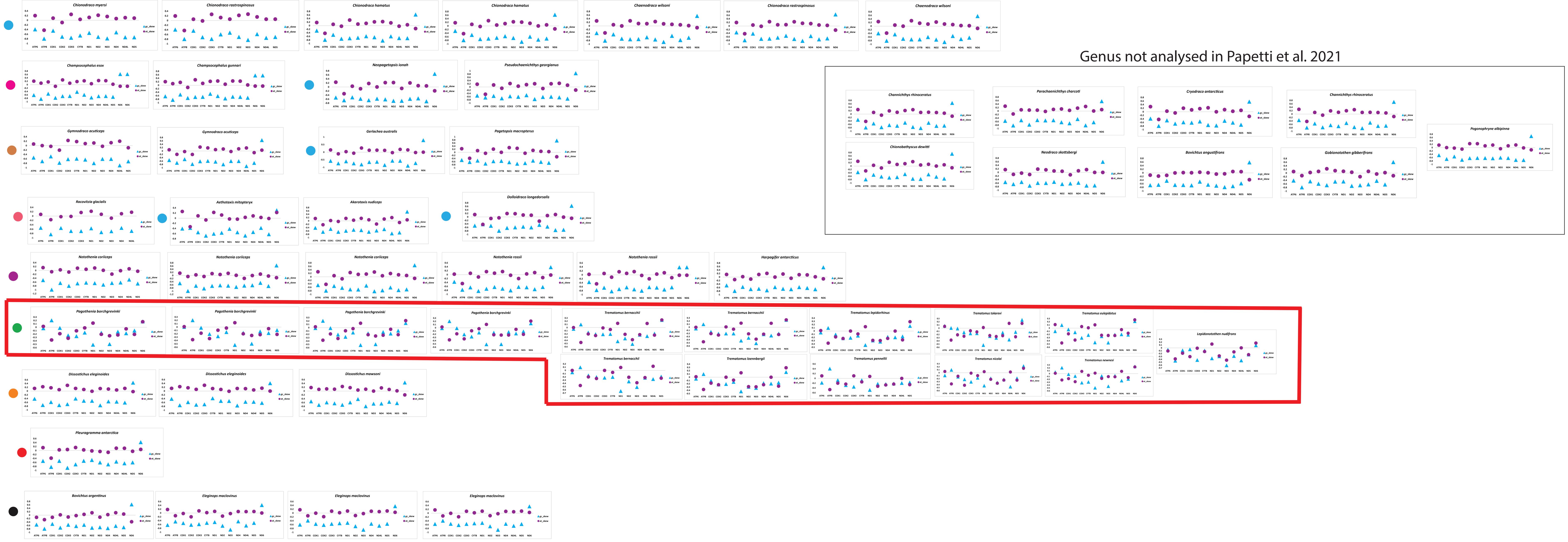

### Fig.S6.jpg

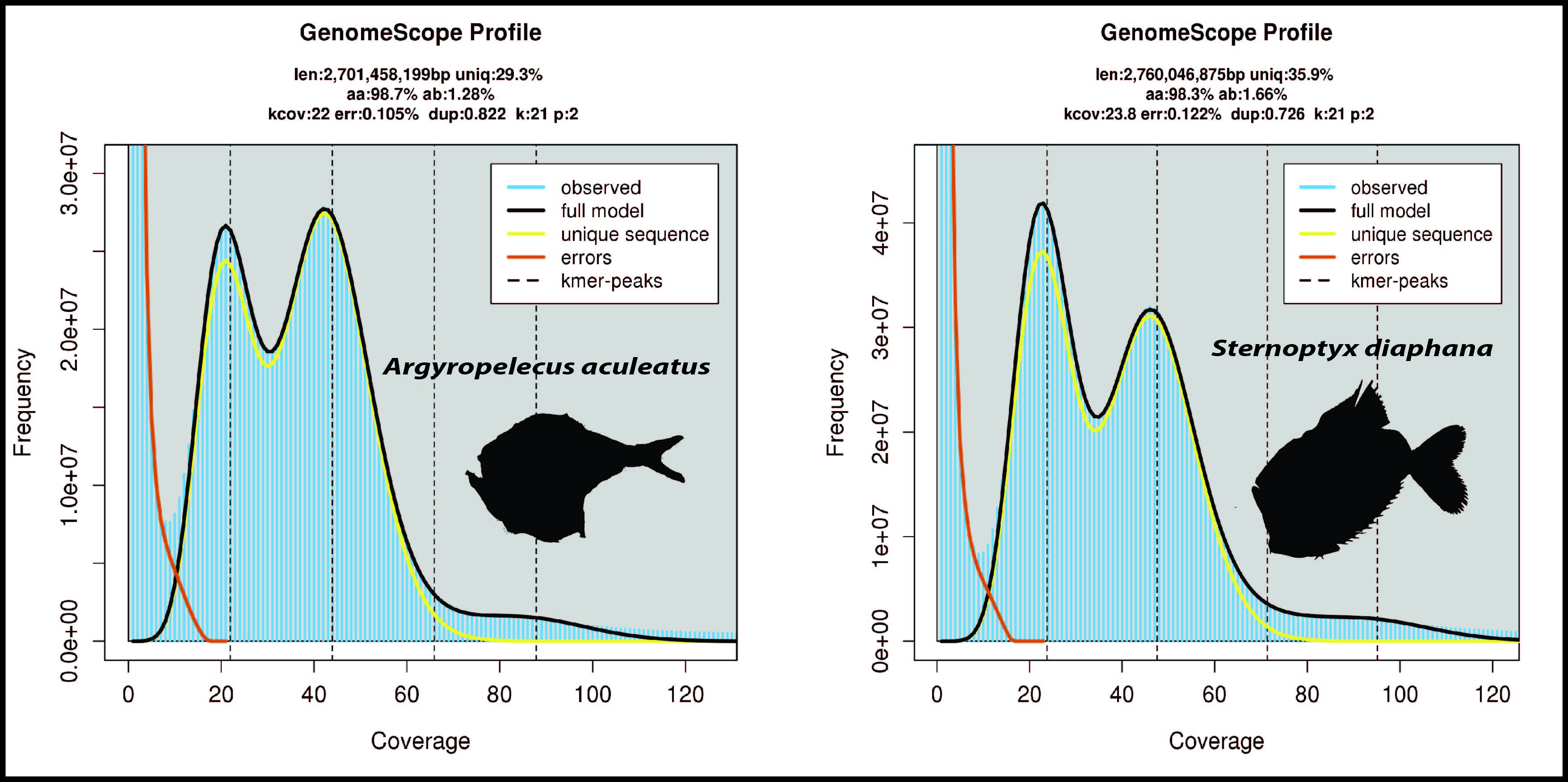

### Fig.S7.jpg

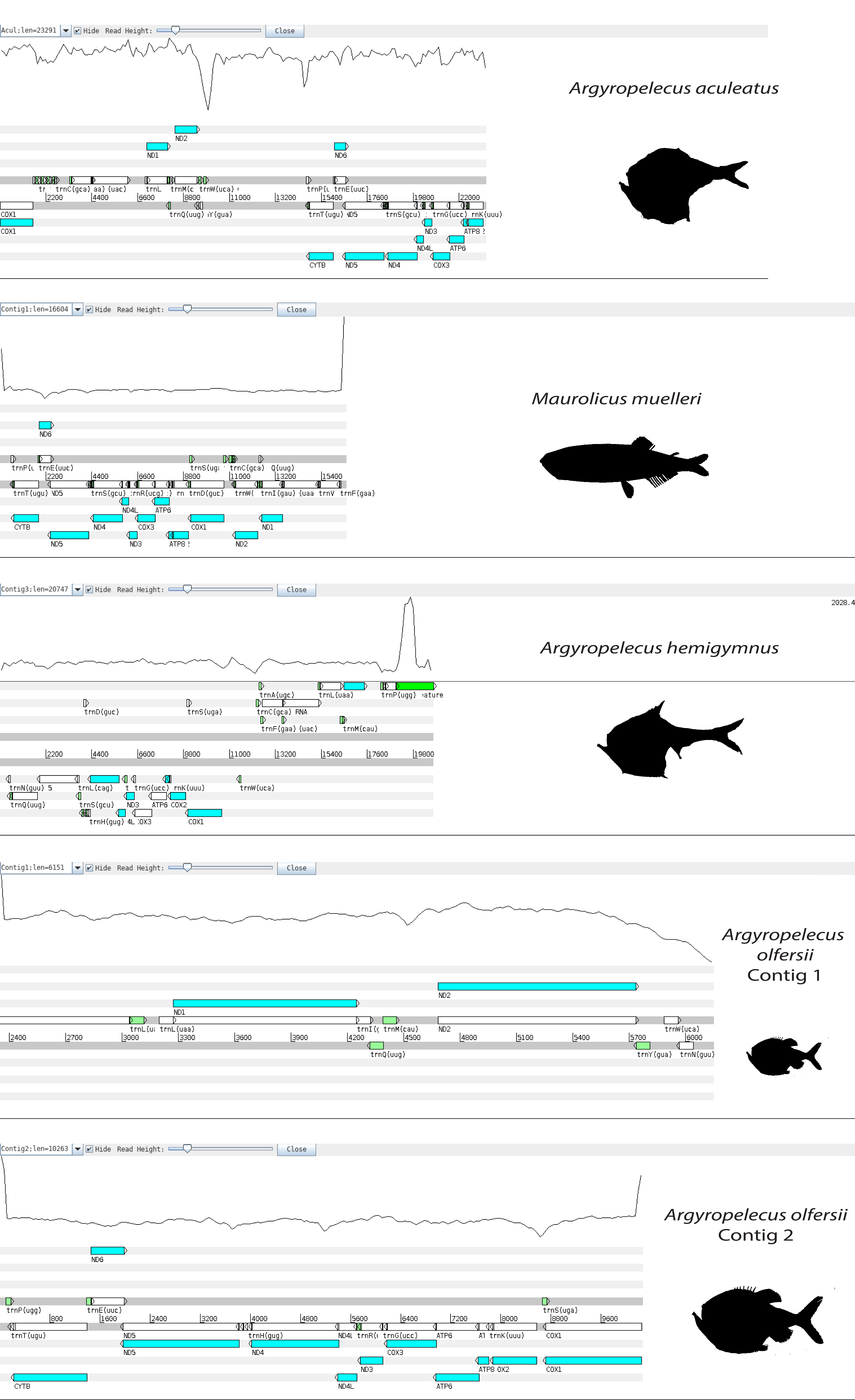
